# Pyruvate Kinase Directly Generates GTP in Glycolysis, Supporting Growth and Contributing to Guanosine Toxicity

**DOI:** 10.1101/2024.12.17.629031

**Authors:** Fukang She, Kuanqing Liu, Brent W Anderson, Tippapha Pisithkul, Yanxiu Li, Danny K. Fung, Tyler McCue, William Mulhern, Daniel Amador-Noguez, Jue D. Wang

## Abstract

Guanosine triphosphate (GTP) is essential for macromolecular biosynthesis, and its intracellular levels are tightly regulated in bacteria. Loss of the alarmone (p)ppGpp disrupts GTP regulation in *Bacillus subtilis*, causing cell death in the presence of exogenous guanosine and underscoring the critical importance of GTP homeostasis. To investigate the basis of guanosine toxicity, we performed a genetic selection for spontaneous mutations that suppress this effect, uncovering an unexpected link between GTP synthesis and glycolysis. In particular, we identified suppressor mutations in *pyk*, which encodes pyruvate kinase, a glycolytic enzyme. Metabolomic analysis revealed that inactivating pyruvate kinase prevents guanosine toxicity by reducing GTP levels. Although traditionally associated with ATP generation via substrate-level phosphorylation, *B. subtilis* pyruvate kinase *in vitro* was found to produce GTP and UTP approximately ten and three times more efficiently than ATP, respectively. This efficient GTP/UTP synthesis extends to *Enterococcus faecalis* and *Listeria monocytogenes*, challenging the conventional understanding of pyruvate kinase’s primary role in ATP production. These findings support a model in which glycolysis directly contributes to GTP synthesis, fueling energy-demanding processes such as protein translation. Finally, we observed a synergistic essentiality of the Δ*ndk* Δ*pyk* double mutant specifically on glucose, indicating that pyruvate kinase and nucleoside diphosphate kinase are the major contributors of NTP production and complement each other during glycolysis. Our work highlights the critical role of nucleotide selectivity in pyruvate kinase and its broader implications in cellular physiology

**Importance:** In this study, we reveal pyruvate kinase, a key glycolytic enzyme, primarily generates GTP from GDP in *Bacillus subtilis*, relatively to other trinucleotides such as ATP. This finding, uncovered through genetic selection for mutants that suppress toxic GTP overaccumulation, challenges the conventional understanding that pyruvate kinase predominantly produces ATP via substrate-level phosphorylation. The substantial role of GTP production by pyruvate kinase suggests a model where glycolysis rapidly and directly supplies GTP as the energy currency to power high GTP-demanding processes such as protein synthesis. Our results underscore the importance of nucleotide selectivity (ATP vs. GTP vs UTP) in shaping the physiological state and fate of the cell, prompting further exploration into the mechanisms and broader implications of this selective nucleotide synthesis.

## Introduction

Purine nucleotides GTP and ATP are crucial small molecules in cell physiology, serving as building blocks for nucleic acid synthesis, participants in signaling and enzymatic reactions, and carriers of cellular energy for metabolism. While ATP is widely recognized as the general energy currency of the cell, GTP specifically powers processes such as protein synthesis (1, 2). Both GTP and ATP are synthesized from the common intermediate IMP, a product of the *de novo* purine synthesis pathway (3). From IMP, ADP and GDP are generated through separate pathways. While ATP is produced from ADP by both substrate-level phosphorylation and oxidative phosphorylation (4), GTP is produced from GDP by the enzyme nucleoside-diphosphate kinase using ATP as the phosphate donor. In addition to the *de novo* pathway, GTP and ATP can also be produced via the salvage pathway (3), which efficiently recycles purine nucleosides and bases from the environment.

The synthesis of GTP is tightly regulated in both the *de novo* and salvage purine biosynthesis pathways (4–6). In bacteria, the alarmones guanosine tetra- and pentaphosphate, collectively known as (p)ppGpp (7), accumulate during nutrient stress and inhibit GTP synthesis, representing a crucial mechanism for maintaining nucleotide homeostasis (8–13). In Gram-positive bacteria, including the soil bacterium *Bacillus subtilis* and human pathogens such as *Enterococcus faecalis* and *Staphylococcus aureus*, (p)ppGpp tunes GTP levels in response to nutritional and other environmental cues to regulate growth, ribosome biogenesis, and translation (14–16). In addition, loss of (p)ppGpp renders these bacteria highly sensitive to exogenous guanosine, leading to drastic loss of vitality even in nutrient-rich conditions (13, 17, 18). This highlights the conserved role of (p)ppGpp in controlling GTP levels and maintaining bacterial survival under varying environmental conditions.

In this study, we conducted a genetic selection in *B. subtilis* to identify mutations that suppress guanosine toxicity in cells lacking (p)ppGpp. Unexpectedly, we repeatedly identified loss-of-function mutations in the *pyk* gene, which encodes pyruvate kinase, a key enzyme in glycolysis. An established function of pyruvate kinase is to catalyze the final step of glycolysis by converting phosphoenolpyruvate (PEP) to pyruvate, concomitantly generating ATP through substrate-level phosphorylation by transferring a phosphate group from PEP to ADP (4). As a key enzyme in energy production, pyruvate kinase is crucial both for ATP generation and fueling the TCA cycle, thus driving cellular respiration and overall energy metabolism (19, 20).

Our findings reveal that pyruvate kinase efficiently generates GTP during substrate-level phosphorylation, with GDP serving as the preferred phosphate acceptor in *B. subtilis* and related Firmicutes. These findings challenge the conventional view that pyruvate kinase primarily produces ATP in the cell. We conclude that pyruvate kinase plays a key role in GTP production in *B. subtilis* and contributes to guanosine toxicity in the absence of (p)ppGpp. Furthermore, we observed that the Δ*ndk* Δ*pyk* double mutant is unable to grow on glucose as the sole carbon source, indicating that pyruvate kinase and nucleoside diphosphate kinase are the primary contributors to NTP production during glycolysis. We discuss the hypothesis that direct and selective production of NTPs via substrate-level phosphorylation is a widespread mechanism in optimizing energy production and resource allocation.

## Results

### Guanosine treatment results in drastic expansion of guanine nucleotide pools and loss of viability in (p)ppGpp^0^ cells

In *B. subtilis*, GTP is produced through *de novo* or salvage pathways that convert acquired guanine or guanosine into guanine nucleotides. (p)ppGpp directly inhibits GTP synthesis by facilitating PurR-mediated transcriptional repression of purine biosynthesis genes, as well as through direct inhibition of key purine synthesis enzymes including IMP dehydrogenase (IMPDH), guanylate kinase (Gmk), and hypoxanthine-guanine phosphoribosyltransferase (HprT) (Fig. 1A) (13, 21). The loss of (p)ppGpp results in pleotropic defects, the most intriguing of which is the disruption of GTP homeostasis and accumulation of abnormally high cellular GTP in the presence of exogenous guanosine (13).

**FIG 1.**
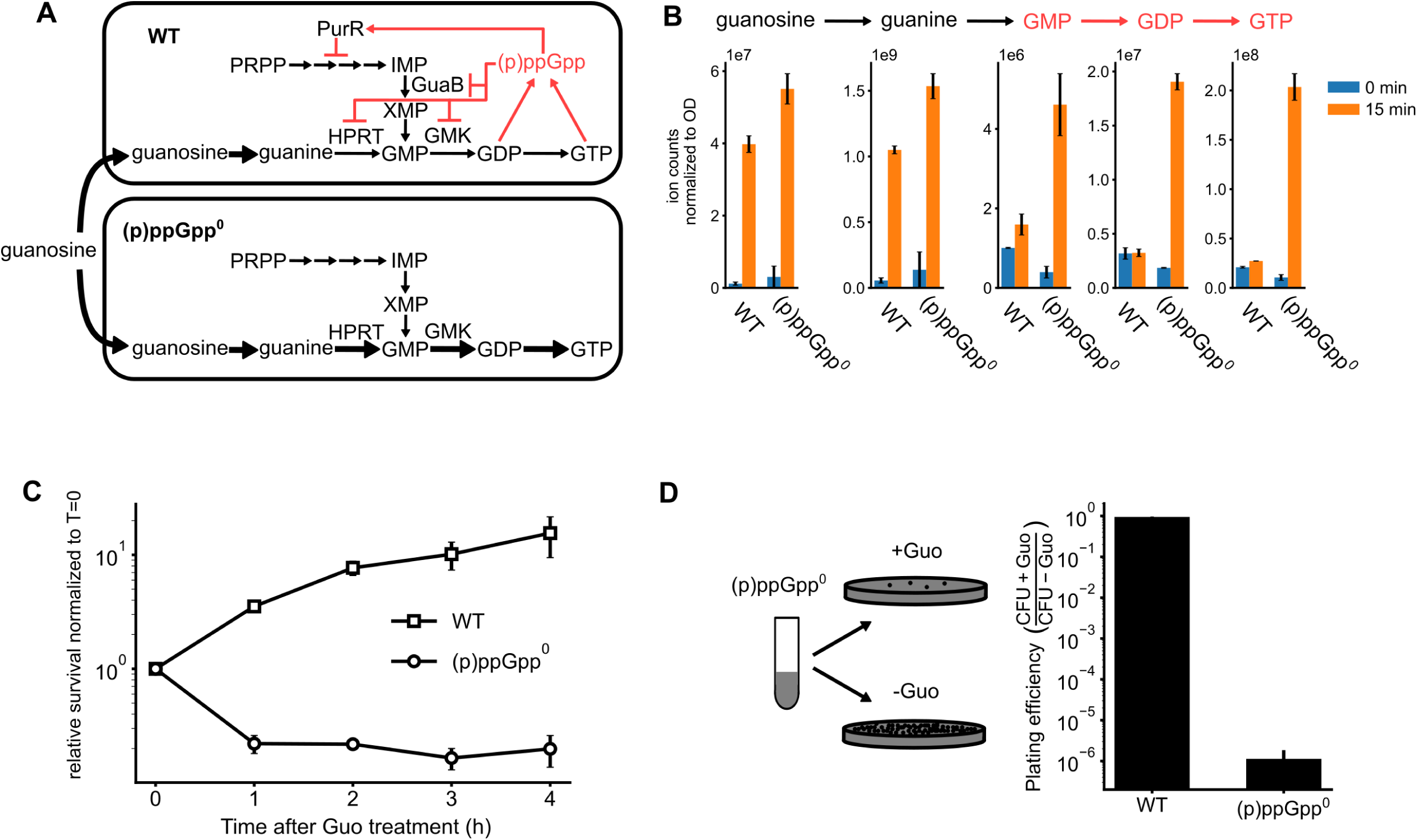
(p)ppGpp^0^ cells are sensitive to extracellular guanosine. (A) Schematics of guanosine utilization and guanine nucleotide synthesis in wild-type (WT) and (p)ppGpp^0^ cells. Both the *de novo* pathway (from PRPP) and the salvage pathway (from extracellular guanosine) of the GTP biosynthesis are negatively regulated by (p)ppGpp (red). (B) The guanine nucleobase, nucleosides and nucleotides levels in WT and (p)ppGpp^0^ cells before guanosine treatment and 15 min after guanosine treatment. Metabolites were measured using LC-MS and normalized to OD_600_. (C) Survival of the wild-type (WT) and (p)ppGpp^0^ cells after 1mM of guanosine is added into defined liquid media supplemented with glucose and CAS-amino acids. (D) Plating efficiency of wild-type (WT) and (p)ppGpp^0^ cells on solid medium supplemented with glucose and CAS-amino acids. CFU: Colony forming units, with (+Guo) or without (-Guo) guanosine. Error bars represent the standard error of the mean from three replicates. Unless otherwise stated.

To track guanosine metabolism in cells lacking (p)ppGpp, we conducted liquid chromatography coupled with mass spectrometry (LC-MS) in *B. subtilis* wild-type and a mutant lacking all three (p)ppGpp synthetases (designated as (p)ppGpp^0^). Following guanosine treatment, both wild-type and (p)ppGpp^0^ cells accumulated guanosine and guanine to similar extent, suggesting that their uptake is comparable between the two strains. However, levels of GMP, GDP, and GTP rose much higher in (p)ppGpp^0^ cells compared to in wild-type cells (Fig. 1B). The overaccumulation of guanine nucleotides in (p)ppGpp^0^ also resulted in rapid cell death in liquid cultures after guanosine treatment (Fig. 1C), as well as inability to form colonies on solid media supplemented with guanosine (Fig. 1D). Overall, the results demonstrate that (p)ppGpp is absolutely necessary to maintain guanosine nucleotide homeostasis in the presence of exogenous guanosine, preventing a potentially toxic buildup of GMP, GDP and GTP.

### Selection for spontaneous suppressors that rescue guanosine toxicity in (p)ppGpp^0^

To investigate the mechanism of guanosine toxicity, we performed an unbiased genetic selection for suppressors that can overcome the inability of (p)ppGpp^0^ cells to form colonies in the presence of exogenous guanosine (Fig. 2A). In this assay, we used a mutant that is completely devoid of all three (p)ppGpp synthetases to prevent generation of suppressors that regain (p)ppGpp synthesis function. Liquid cultures of (p)ppGpp^0^ cells were plated on selective media containing glucose, amino acids, and guanosine. While these cells could not form colonies due to guanosine toxicity, rare suppressor mutants occasionally emerged. These mutants were further confirmed for their ability to grow on guanosine agar plates (Fig. 2B, C and D), followed by detailed measurement of their GTP levels after guanosine treatment (Fig. 2E, F and G), and their growth rates in liquid media (Fig. 2H, I and J). Subsequent DNA sequencing revealed mutations at three genetic loci: *gmk*, *hprT* and *pyk*, encoding guanylate kinase (Gmk), hypoxanthine-guanine phosphotransferase (HprT), and pyruvate kinase (PK), respectively.

**FIG 2.**
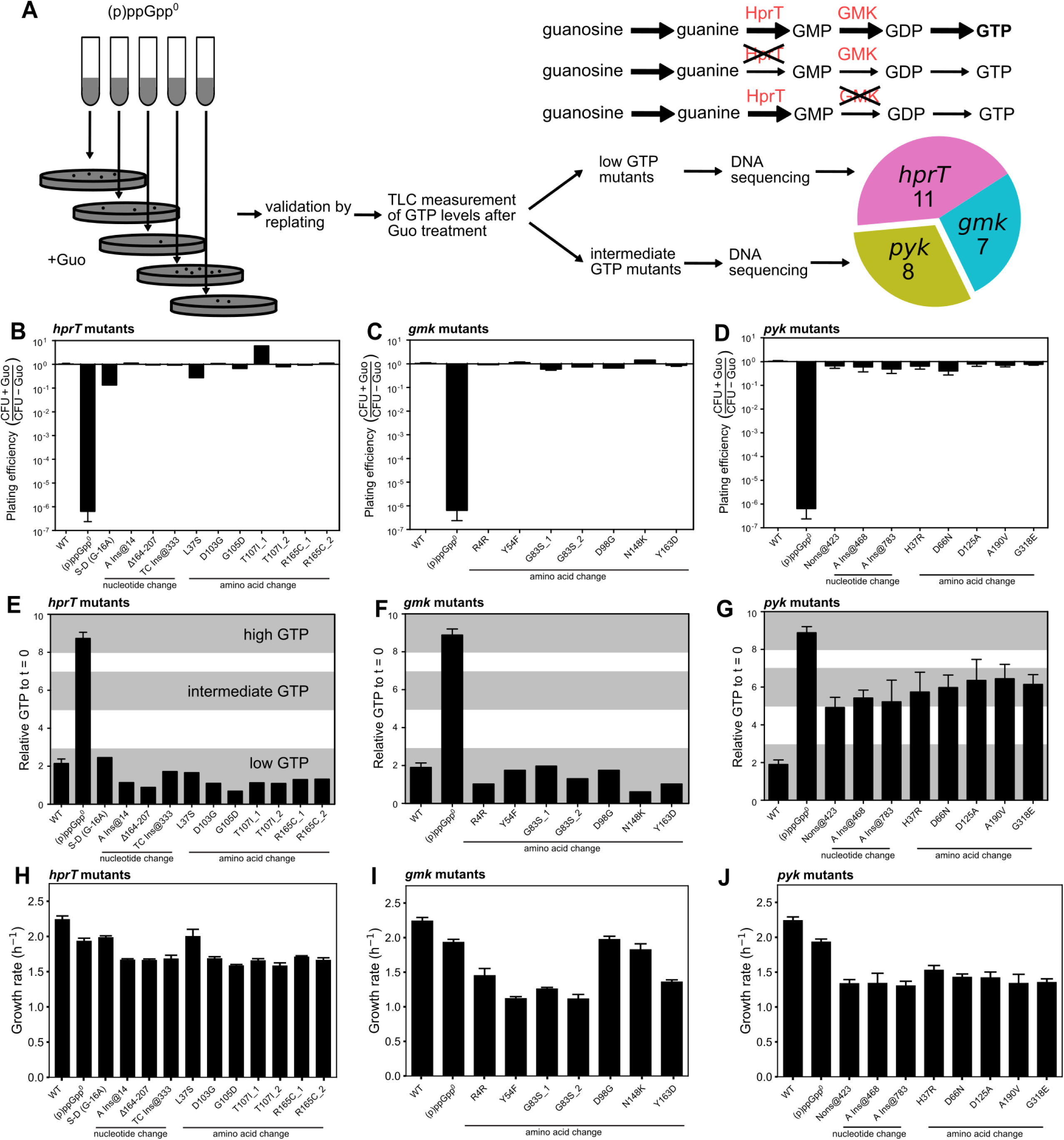
Genetic selection and characterization of suppressors of guanosine toxicity in (p)ppGpp^0^ *B. subtilis* cells. (A) Schematics of suppressor selection and characterization. Briefly, (p)ppGpp^0^ cells growing in independent liquid cultures were plated on solid media supplemented with 0.1 mM guanosine, one suppressor from each plate was isolated. Mutants were whole genome-sequenced or Sanger-sequenced to identify the associated suppressor mutations. 11 *hprT*, 7 *gmk* and 8 *pyk* suppressor mutations were identified. *hprT* and *gmk* mutations would block the conversion of guanosine to GTP at different steps. (B-D): Plating efficiency of *hprT* mutants (B), *gmk* mutants (C), and *pyk* mutants (D) was determined by dividing colony forming units (CFUs) of the same culture plated on agar plates with or without guanosine. (E-G): Changes in GTP levels in *hprT* mutants (E), *gmk* mutants (F), and *pyk* mutants (G) following guanosine treatment were measured by thin layer chromatography (TLC). (H-J): Growth rates of *hprT* mutants (H), *gmk* mutants (I), and *pyk* mutants (J) in liquid media without guanosine. Annotation for mutations: S-D: Shine-Dalgarno sequence; Ins: insertion; Δ: deletion. The *hprT* T107I_1 and _2 mutants and R165C_1 and _2 mutants, as well as the *gmk* G83S_1 and _2 mutants, were identified from independent liquid cultures. *gmk* R4R mutant is a synonymous mutation from AGA to AGG.

We found that suppressors robustly grow on guanosine plates despite lacking (p)ppGpp (Figs. 2B, C and D). GTP levels varied among suppressors, which *hprT* mutants and *gmk* mutants exhibiting “low GTP”: GTP levels similar to or lower than wild-type cells (Figs. 2E and 2F), and *pyk* mutants exhibiting “intermediate GTP”: GTP levels lower than (p)ppGpp^0^ but significantly higher than wild-type cells (Fig. 2G). None of the suppressors had GTP levels as high as those of (p)ppGpp^0^ cells.

The genes *hprT* and *gmk* encode HprT and Gmk, two enzymes responsible for producing GMP from guanine and GDP from GMP, respectively (Fig. 2A). Both enzymes are also known to be strongly inhibited by (p)ppGpp to regulate GTP levels in wild-type cells (13). As inactivation of either enzyme would be expected to curb conversion of guanosine to guanine nucleotides, we first performed Sanger sequencing at *hprT* and *gmk* loci in “low GTP” suppressors and indeed identified 11 *hprT* and 7 *gmk* suppressor mutations. The *hprT* suppressors exhibited insertions, deletions, or base substitutions that altered either amino acids or the putative Shine-Dalgarno sequence, which together with their hampered GTP accumulation in the presence of guanosine (Fig. 2B), suggest that they are loss-of-function mutations. Although *gmk* suppressors only had base-substitutions in coding sequences likely because *gmk* is an essential gene and its complete inactivation would be lethal, they are nonetheless loss-of-function mutations based on the fact all suppressor mutations displayed dramatically lower GTP levels than (p)ppGpp^0^ cells following guanosine treatment (Fig. 2C). The nonsynonymous substitutions in *gmk* are located in residues near the active site of the enzyme (22). In addition, a synonymous substitution (AGA to AGG at codon 4, both encoding arginine) in *gmk* is likely to result in lower expression of Gmk, as AGG is a rare codon compared to AGA (23). The *gmk* mutations identified herein are similar to those uncovered in our previous genetic selection under amino acid starvation (13), which had lower GDP and GTP but high GMP due to reduced Gmk activity. Since *gmk* mutants are resistant to guanosine while having high GMP and other upstream intermediates, it is likely that guanosine toxicity is largely due to high GDP or GTP levels. We conclude that the *hprT* and *gmk* suppressors relieve guanosine toxicity due to their loss-of-function mutations, mimicking (p)ppGpp’s ability to inhibit HprT and Gmk activity (Fig. 2A).

### Loss-of-function mutations in *pyk* rescued guanosine toxicity

Intriguingly, eight suppressors with substantial growth and intermediate GTP levels after guanosine treatment did not have any mutations in *hprT* or *gmk*, suggesting a novel suppression route. Indeed, subsequent whole genome and Sanger sequencing revealed that they all contained mutations in *pyk*, which encodes the glycolytic enzyme pyruvate kinase (Table 1). Pyruvate kinase catalyzes the conversion of PEP to pyruvate, feeding into the TCA cycle (Fig. 3A). The mutations of the eight *pyk* suppressors include frameshift insertions, nonsense mutations, and missense mutations (Table 1). Based off the sequence alignment of pyruvate kinase homologs and the crystal structure of *G. stearothermophilus* pyruvate kinase (PDB ID: 2E28) (24), all residues corresponding to the identified missense mutations are conserved and located near the active site (Fig. 3B, Fig. S1), prompting us to speculate that they would likely result in loss-of-function of pyruvate kinase.

**FIG 3.**
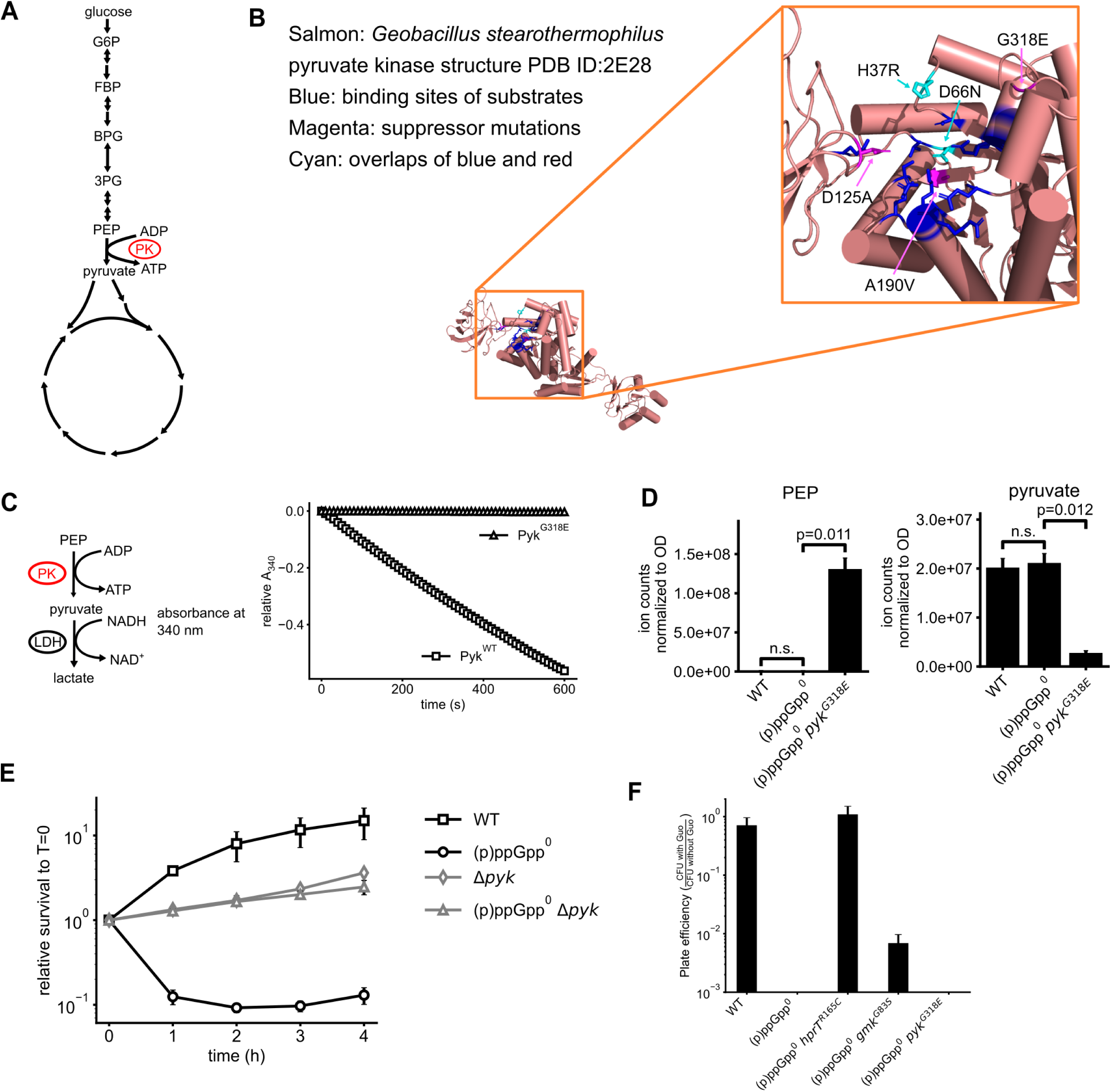
Loss-of-function mutations in *pyk* rescue guanosine toxicity of (p)ppGpp^0^ cells. (A) Schematics of glycolysis and the tricarboxylic acid (TCA) cycle. Pyruvate kinase catalyzes the last step of glycolysis, converting PEP and ADP to pyruvate and ATP. G6P: glucose 6-phosphate; FBP: fructose 1,6-bisphosphate; BPG: 1,3-bisphosphoglycerate; 3PG: 3-phosphoglycerate; PEP: phosphoenolpyruvate; PK: pyruvate kinase. (B) *pyk* suppressor missense mutations mapped to the structure of *G. stearothermophilus* pyruvate kinase (PDB ID: 2E28). Altered amino acids were all located near the active site. (C) Pyruvate kinase variant with the suppressor mutation G318E is catalytically inactive *in vitro*. Pyruvate kinase activity was monitored by pyruvate production, which was subsequently converted to lactate by lactate dehydrogenase (LDH) in the presence of NADH. This reaction resulted in a decrease in absorbance at 340 nm, corresponding to NADH consumption. (D) Intracellular PEP and pyruvate levels in WT, (p)ppGpp^0^ and (p)ppGpp^0^ *pyk^G318E^* cells measured by LC-MS. The accumulation of PEP and diminishment of pyruvate indicates the loss of pyruvate kinase activity. (E) Survival of WT, (p)ppGpp^0^, Δ*pyk* and (p)ppGpp^0^ Δ*pyk* cells in a time course of 1 mM guanosine treatment. (F) Plating efficiency of WT, (p)ppGpp^0^, and representative *hprT*, *gmk* and *pyk* suppressor mutations on plates supplemented with malate instead of glucose. CFU with verses without guanosine ratio is plotted.

**TABLE 1.** Suppressors of (p)ppGpp^0^ guanosine toxicity.

| genes | Mutations (nt) | Type | Plating efficiency<sup>a</sup> | GTP level<sup>b</sup> | strain |
| --- | --- | --- | --- | --- | --- |
| *hprT* | -16 G>A | RBS mutation | 1e-1 | 2.2 | JDW1854 |
| *hprT* | +14 A insertion | frameshift | 1 | 1.1 | JDW1850 |
| *hprT* | *hprT*<sup>L37S</sup> | Leu37Ser | 3e-1 | 1.3 | JDW1839 |
| *hprT* | 164-207 deletion | Deletion | 9e-1 | 1.0 | JDW1853 |
| *hprT* | *hprT*<sup>D103G</sup> | Asp103Gly | 1 | 1.1 | JDW1848 |
| *hprT* | *hprT*<sup>G105D</sup> | Gly105Asp | 8e-1 | 0.7 | JDW1858 |
| *hprT* | *hprT*<sup>T107I</sup> | Thr107Ile | 7e-1 | 1.2 | JDW1852 |
| *hprT* | *hprT*<sup>T107I</sup> | Thr107Ile | 6 | 1.1 | JDW1856 |
| *hprT* | *hprT*<sup>R165C</sup> | Arg165Cys | 9e-1 | 1.4 | JDW1849 |
| *hprT* | *hprT*<sup>R165C</sup> | Arg165Cys | 1 | 1.4 | JDW1851 |
| *hprT* | +333 TC insertion | Insertion | 9e-1 | 1.9 | JDW1834 |
| *gmk* | *gmk*<sup>R4R</sup> | nucleotide A12G | 9e-1 | 1.2 | JDW1835 |
| *gmk* | *gmk*<sup>Y54F</sup> | Tyr54Phe | 1 | 1.6 | JDW1837 |
| *gmk* | *gmk*<sup>G83S</sup> | Gly83Ser | 5e-1 | 2.1 | JDW1841 |
| *gmk* | *gmk*<sup>G83S</sup> | Gly83Ser | 7e-1 | 1.2 | JDW1842 |
| *gmk* | *gmk*<sup>D98G</sup> | Asp98Gly | 7e-1 | 1.6 | JDW1833 |
| *gmk* | *gmk*<sup>N148K</sup> | Asn148Lys | 1 | 0.6 | JDW1826 |
| *gmk* | *gmk*<sup>Y163D</sup> | Tyr163Asp | 1 | 1.0 | JDW1836 |
| *pyk* | *pyk*<sup>H37R</sup> | His37Arg | 3e-1 | 6.8±3.9 | JDW1847 |
| *pyk* | *pyk*<sup>D66N</sup> | Asp66Asn | 2e-1 | 5.8±0.9 | JDW1843 |
| *pyk* | *pyk*<sup>D125A</sup> | Asp125Ala | 6e-1 | 5.7±0.2 | JDW1838 |
| *pyk* | *pyk*<sup>A190V</sup> | Ala190Val | 7e-1 | 4.6±0.2 | JDW1840 |
| *pyk* | *pyk*<sup>G318E</sup> | Gly318Glu | 7e-1 | 6.1±1.6 | JDW1857 |
| *pyk* | +423 Nonsense | Nonsense | 4e-1 | 5.1±0.8 | JDW1832 |
| *pyk* | +468 A insertion | frameshift | 2e-1 | 4.6±0.4 | JDW1846 |
| *pyk* | +783 A insertion | frameshift | 4e-1 | 4.2 | JDW1830 |

a. CFU on Min-CAS-Guo/CFU on Min-CAS
b. GTP levels normalized to GTP levels at T=0 plus/minus standard error

To experimentally test our hypothesis, we recombinantly expressed and purified wild-type *B. subtilis* pyruvate kinase and a G318E mutant we identified from the suppressor selection and performed enzymatic assays using a coupled reaction comprising pyruvate kinase and lactate dehydrogenase (LDH) (25). Wild type pyruvate kinase could rapidly convert PEP to pyruvate, but we could barely detect any activity in the G318E mutant (Fig. 3C), confirming that it is indeed a loss-of-function mutation. Consistent with our *in vitro* results, LC-MS based metabolomics showed that PEP (upstream of pyruvate kinase) was strongly elevated in the *pyk^G318E^* mutant, while pyruvate (its downstream product) was significantly depleted (Fig. 3D). Although we couldn’t generate a strain to complement the *pyk^G318E^* mutant with a wild-type copy to prove that the point mutants are loss-of-function due to the loss of transformability in (p)ppGpp^0^ mutant, we bypassed it by making a deletion of *pyk* in *B. subtilis* and introducing subsequent deletions of (p)ppGpp synthetase genes *sasA*, *sasB*, and *relA*. The resulting (p)ppGpp^0^ Δ*pyk* mutant is resistant to guanosine (Fig. 3E), suggesting that inactivation of *pyk* is sufficient to rescue the guanosine toxicity. Finally, because PK is a glycolytic enzyme, we examined whether its rescue of guanosine toxicity requires glucose. We found that in the absence of glucose, when grown on malate, (p)ppGpp^0^ *pyk* mutant could no longer suppress guanosine toxicity, although (p)ppGpp^0^ *hprT* and *gmk* mutants still remained resistant to guanosine (Fig. 3F). Collectively, these results support the notion that pyruvate kinase activity during glycolysis contributes to guanosine toxicity, and inactivation of pyruvate kinase rescues guanosine toxicity.

### Rescue of guanosine toxicity by pyruvate kinase inactivation is unlikely due to changes in central carbon metabolism

To explore how pyruvate kinase inactivation might suppress guanosine toxicity, we conducted LC-MS-based metabolomic profiling of wild type, (p)ppGpp^0^, and (p)ppGpp^0^ *pyk*^G318E^ strains before and after guanosine treatment, focusing on key metabolites in central metabolism (Fig. S2). We chose to focus on the (p)ppGpp^0^ *pyk*^G318E^ strain for the following reasons: Firstly, It displayed one of the most robust growth rescue phenotypes for *pyk* suppressors; Secondly, It still harbors an intact albeit mutated *pyk*, which we believe causes significantly less perturbation to normal cellular physiology compared to complete *pyk* deletion, as many of the glycolytic enzymes may form transient multi-enzyme assemblies (26) and many may have moonlighting functions as well (27); Thirdly, We also confirmed that it did not carry any secondary mutations by whole genome sequencing.

We first examined the hypothesis that pyruvate kinase inactivation rescues guanosine toxicity by altering carbon metabolism. To test this idea, we first analyzed metabolites downstream of pyruvate kinase, including pyruvate and TCA cycle intermediates. As expected, the *pyk ^G318E^* suppressor showed reduced levels of pyruvate and TCA intermediates than (p)ppGpp^0^ cells (Fig. 4A). However, in the (p)ppGpp^0^ cells, these metabolites were only mildly elevated compared to wild-type cells and were reduced upon guanosine treatment, indicating that their accumulation is not the cause of toxicity (Fig. 4A). Furthermore, supplementing (p)ppGpp^0^ *pyk* mutants with glucose and pyruvate did not restore toxicity (Fig. S3), confirming that *pyk* mutants did not rescue guanosine toxicity by depleting pyruvate pool or TCA cycle.

**FIG 4.**
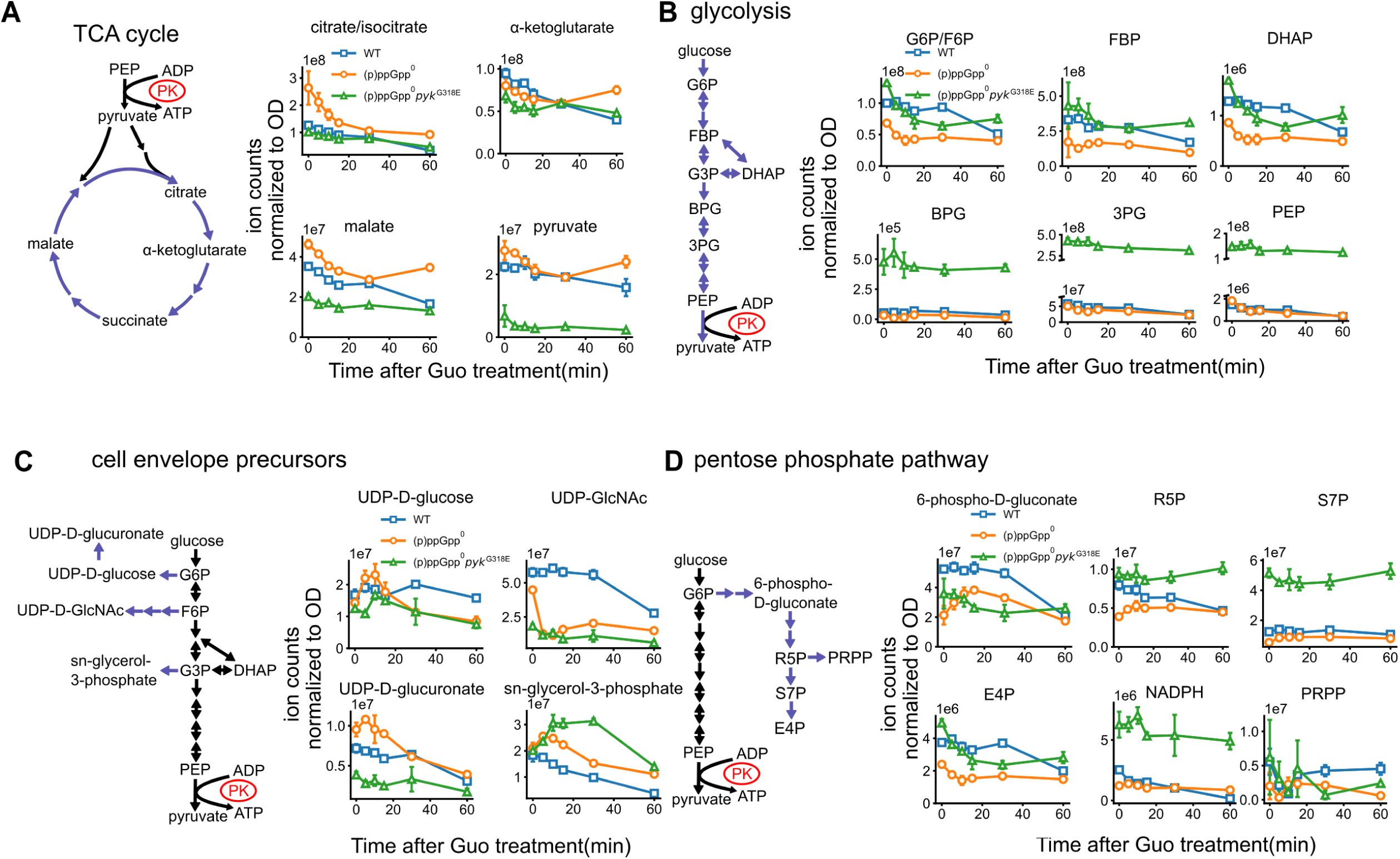
Central carbon metabolism after guanosine treatment in WT, (p)ppGpp^0^ and (p)ppGpp^0^ *pyk^G318E^* cells. Metabolites were quantified by LC-MS and plotted separately for TCA cycle (A), glycolysis (B), cell envelope precursors (C) and the pentose phosphate pathway (D), in a time course of 1 mM guanosine treatment. LC-MS signals are normalized to OD_600_. Guo: guanosine; G6P/F6P: glucose 6-phosphate or fructose 6-phosphate; FBP: fructose 1,6-bisphosphate; G3P: glyceraldehyde 3-phosphate; DHAP: dihydroxyacetone phosphate; BPG: 1,3-bisphosphoglycerate; 3PG: 3-phosphoglycerate; PEP: phosphoenolpyruvate; UDP-GlcNAc: UDP N-acetylglucosamine; R5P: ribose 5-phosphate; S7P: sedoheptulose 7-phosphate; E4P: erythrose 4-phosphate; PRPP: 5-phosphoribosyl 1-pyrophosphate.

We next tested the idea that pyruvate kinase inactivation conserves glycolytic intermediates to allow appropriate resource allocation (8, 28). We reasoned that in the absence of (p)ppGpp, exogenous guanosine triggers excessive guanine nucleotide production, which can deplete glycolytic intermediates due to increased flux into the pentose phosphate pathway. This misdirection of resources can disrupt the synthesis of other essential building blocks derived from glycolytic intermediates, such as peptidoglycan and fatty acids. By inactivating pyruvate kinase, the glycolytic intermediate phosphoenolpyruvate (PEP) is prevented from being converted to pyruvate and subsequently entering the TCA cycle. This effectively conserves glycolytic intermediates for anabolic processes.

To test this model, we examined glycolytic intermediates upstream of pyruvate kinase, from glucose 6-phosphate (G6P) to PEP (Fig 4B). In (p)ppGpp^0^ cells, the levels of these metabolites were reduced by ∼2 fold or remained similarly to wild-type levels. The (p)ppGpp^0^ *pyk* mutant display strong elevation of glycolytic intermediates such as PEP, BPG and 3-phosphoglycerate (3PG). Based on the observed reduction of G6P, we considered that pathways relying on G6P, such as the pentose phosphate pathway and cell envelope biogenesis, could be affected. Therefore, we evaluated key biosynthetic precursors from glycolytic intermediates, including those involved in cell envelope biogenesis (Fig. 4C) and the pentose phosphate pathway (Fig. 4D). For cell envelope precursors such as UDP-N-acetylglucosamine (UDP-GlcNAc) and UDP-glucose, there was either no depletion in (p)ppGpp^0^ after guanosine treatment, or the *pyk* mutation did not rescue this depletion. In the pentose phosphate pathway, we observed lower ribose 5-phosphate (R5P) levels in (p)ppGpp^0^, while the *pyk* mutant had higher R5P levels prior to treatment. However, R5P levels increased in (p)ppGpp^0^ after guanosine treatment, suggesting that its depletion is not the cause of toxicity. Additionally, the nucleotide precursor PRPP, a pentose phosphate pathway product, remained largely unaffected (Fig. 4D).

Taken together, our metabolomic analyses suggest that changes in central carbon metabolites, such as PEP and pyruvate, are not the primary cause of guanosine toxicity or growth rescue in the *pyk* mutant.

### The synthesis of GTP from GDP is significantly reduced in the *pyk* suppressor during guanosine treatment

During catalytic conversion of phosphoenolpyruvate (PEP) to pyruvate, pyruvate kinase also transfers a phosphate group from PEP to ADP to produce ATP. Thus, its activity not only plays a key role in carbon metabolism but also contributes to substrate-level phosphorylation to generate NTPs. Since the effect of pyruvate kinase on carbon metabolism has not been shown to be important for the toxicity, we next examined whether its effect on nucleotide synthesis is involved.

We examined the levels of ATP and ADP, as well as other purine and pyrimidine nucleotides to assess potential changes in energy status associated with guanosine toxicity suppression (Fig. 5A, Fig. S4). In agreement with our previous finding (13), guanosine treatment led to a strong accumulation of GTP and a reduction in ATP levels in (p)ppGpp^0^ compared to in wild-type cells. On the other hand, the *pyk* suppressor had reduced GTP and UTP levels by ∼50% while ATP and CTP levels were unaffected (Figs 5B and 5D). In addition, the *pyk* suppressor also had strongly increased GDP levels, even more pronounced than those seen with the (p)ppGpp^0^ cells (Fig. 5D), suggesting that the conversion of GDP to GTP is strongly impeded. Similarly, UDP accumulates in (p)ppGpp^0^ *pyk^G318E^*, suggesting that the conversion of UDP to UTP were also impeded, while conversion to ATP or CTP were less affected (Figs. 5A). This prompts us to speculate that pyruvate kinase might efficiently converts GDP to GTP during guanosine treatment to contribute to death-by-GTP. The loss-of-function mutation in *pyk* could block this conversion, thereby reducing guanosine toxicity.

**FIG 5.**
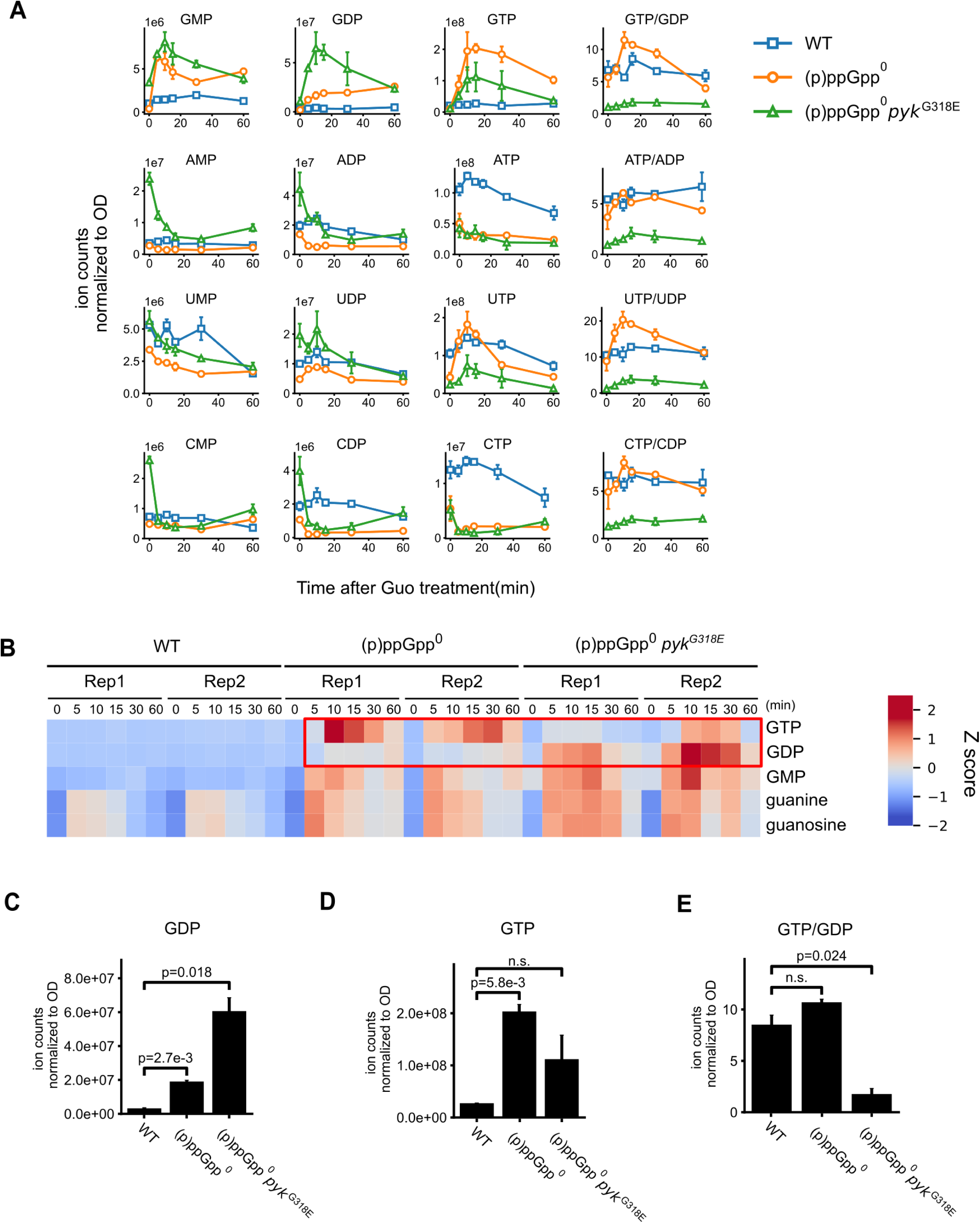
Inactivation of pyruvate kinase impedes conversion of GDP to GTP in (p)ppGpp^0^ cells during guanosine treatment. (A) Nucleotides levels in a time course of guanosine (Guo) treatment in WT, (p)ppGpp^0^ and (p)ppGpp^0^ *pyk^G318E^*, measured using LC-MS and normalized to OD_600_. (B) Heat map showing Z scores of guanine, guanine nucleoside, and guanine nucleotides levels. (C-E) Intracellular levels of GDP (C), GTP (D), and GDP/GTP ratios (E) in WT, (p)ppGpp^0^ and (p)ppGpp^0^ *pyk^G318E^* cells 15 minutes after guanosine treatment.

### *In Vitro* Kinetics Reveal Pyruvate Kinase’s Preference for GTP Synthesis in *Bacillus subtilis* and Other Firmicutes

Pyruvate kinase is conventionally known to produce ATP from ADP during glycolysis (4). The observed reduction in GTP levels in the absence of functional pyruvate kinase could be an indirect result of lower ATP levels, which would in turn affect GTP synthesis by Ndk, the enzyme responsible for converting NDP to NTP using ATP as phosphate donor. However, another possibility is that pyruvate kinase directly plays a role in GTP synthesis. Early *in vitro* studies have shown that pyruvate kinases from *E. coli* and rabbit muscle can use various nucleoside diphosphates as phosphoryl group acceptors (29–31), suggesting that the enzyme might directly produce GTP during substrate level phosphorylation.

To test this hypothesis, we assessed the nucleotide specificity of *B. subtilis* pyruvate kinase *in vitro*. We added ADP, GDP, UDP or CDP individually to reactions containing PEP and monitored pyruvate formation (Figs. 6A, B, C and D). In the absence of other nucleotides, pyruvate kinase activity was higher with ADP compared to other NDP substrates. However, when AMP, a known activator of pyruvate kinase, was added, we observed a marked increase in activity with GDP, UDP and CDP as substrates, but not as much with ADP (Fig. 6A, B, C and D). These findings demonstrate that *B. subtilis* pyruvate kinase can use a variety of nucleoside diphosphates as phosphate acceptors.

**FIG 6.**
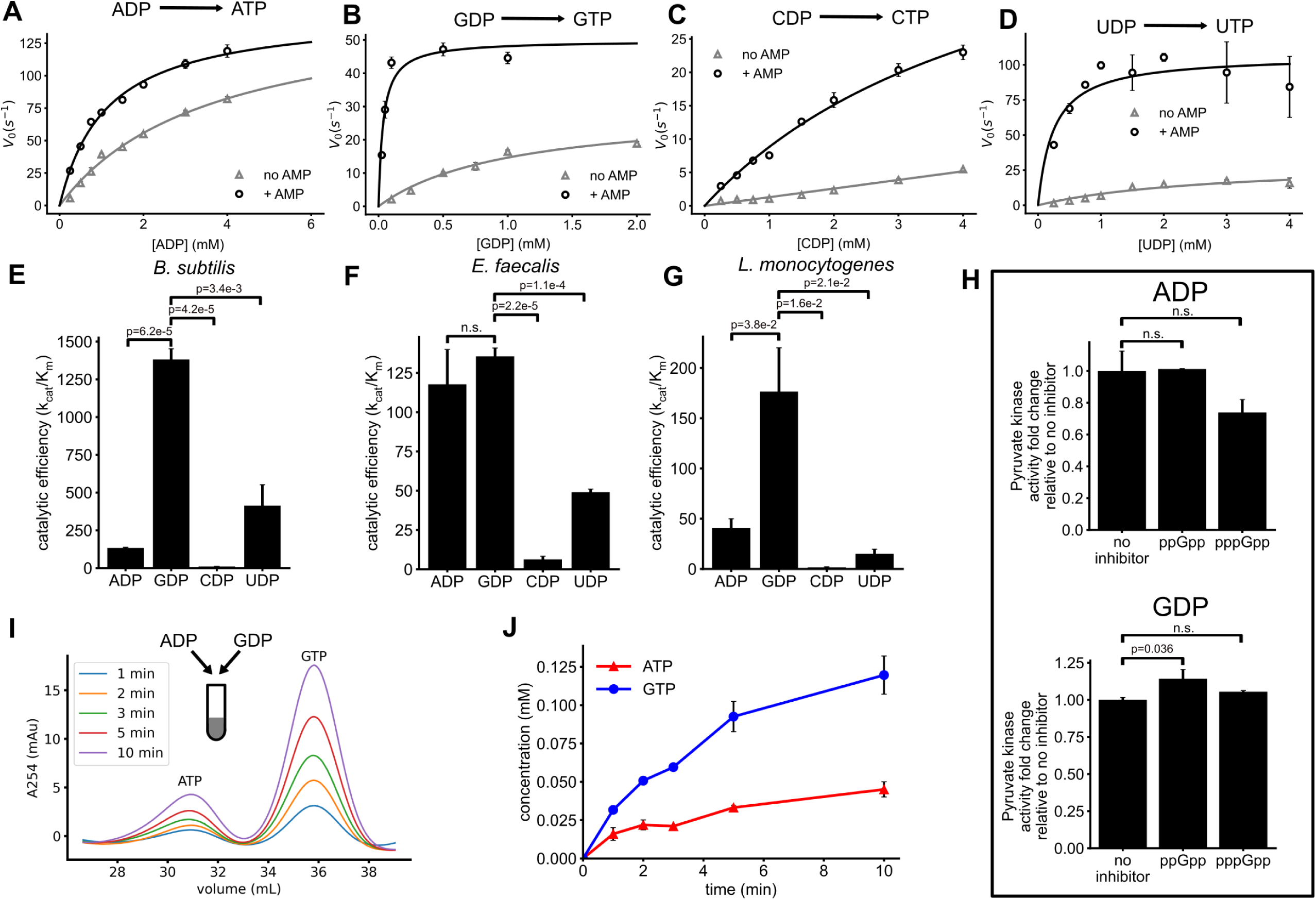
*In vitro* enzyme activity of pyruvate kinases from *B. subtilis* and related Firmicutes using ADP, GDP, CDP or UDP as the substrate. (A-D) *B. subtilis* pyruvate kinase activity (measured by initial velocity) against increasing concentrations of NDP substrates: ADP (A), GDP (B), CDP (C), or UDP (D), with, or without 1 mM AMP as the activator. (E-G) Catalytic efficiencies (k_cat_/K_m_) of pyruvate kinases from *B. subtilis* (E), *E. faecalis* (F), and *L. monocytogenes* (G) using different NDPs as substrates. (H) Effect of 1 mM ppGpp or pppGpp on pyruvate kinase activity using ADP or GDP as the substrate. (I-J) The kinetics of NTP synthesis by pyruvate kinase with PEP and a mixture of 0.5 mM ADP and 0.2 mM GDP as substrates. ATP and GTP products were separated with an anion exchange column and quantified by measuring absorbance at 254 nm. (I) Representative chromatographs at indicated times after the start of reaction. (J) Concentrations of ATP and GTP products over time. Means and standard errors of the mean are obtained from three independent reactions.

We next measured kinetic parameters (K_m_, k_cat_) of *B. subtilis* pyruvate kinase for each NDP in the presence of AMP and compared their catalytic efficiencies (k_cat_/K_m_) (Table 2) (32). Remarkably, GDP emerged as the most preferred substrate, showing a catalytic efficiency 10 times greater than that for ADP (Fig. 6E). UDP was also a favored substrate, with a catalytic efficiency 3 times that of ADP. This correlates with our *in vivo* results (Fig. 5A) where inactivation of pyruvate kinase blocked both GDP-to-GTP and UDP-to-UTP conversions in the cell.

**TABLE 2.** Kinetic parameters of *B. subtilis* pyruvate kinase using different NDPs, +/− 95% confidence intervals. kcat/Km: catalytic efficiencies.

| | $k_{cat}$ ($s^{-1}$) | $K_m$ (mM) | $k_{cat} / K_m$ ($s^{-1} mM^{-1}$) |
| --- | --- | --- | --- |
| ADP | 150±10 | 1.1±0.1 | 134±5 |
| GDP | 50±2 | 0.036±0.001 | 1382±87 |
| CDP | 57±5 | 5.6±0.7 | 10.3±0.5 |
| UDP | 115±15 | 0.35±0.14 | 414±170 |

**TABLE 3.** Strains and Plasmids used in this study.

| Strain or plasmid Name | Genotype | Reference |
| --- | --- | --- |
| YB886 | *trpC2, metB10, xin-1, SPβ* | Yasbin et al. (48) |
| JDW755 | YB886 $\Delta yjbM ywaC::kan relA::mls$ | Kriel et al. (13) |
| JDW1854 | JDW755 *hprT* -16 G>A | This work |
| JDW1850 | JDW755 *hprT* +14 A insertion | This work |
| JDW1839 | JDW755 *hprT*<sup>L37S</sup> | This work |
| JDW1853 | JDW755 *hprT* +164 to +207 deletion | This work |
| JDW1848 | JDW755 *hprT*<sup>D103G</sup> | This work |
| JDW1858 | JDW755 *hprT*<sup>G105D</sup> | This work |
| JDW1852 | JDW755 *hprT*<sup>T107I</sup> | This work |
| JDW1856 | JDW755 *hprT*<sup>T107I</sup> | This work |
| JDW1849 | JDW755 *hprT*<sup>R165C</sup> | This work |
| JDW1851 | JDW755 *hprT*<sup>R165C</sup> | This work |
| JDW1834 | JDW755 *hprT* +333 TC insertion | This work |
| JDW1835 | JDW755 *gmk*<sup>R4R</sup> | This work |
| JDW1837 | JDW755 *gmk*<sup>Y54F</sup> | This work |
| JDW1845 | JDW755 *gmk*<sup>G83S</sup> | This work |
| JDW1841 | JDW755 *gmk*<sup>G83S</sup> | This work |
| JDW1833 | JDW755 *gmk*<sup>D98G</sup> | This work |
| JDW1826 | JDW755 *gmk*<sup>N148K</sup> | This work |
| JDW1836 | JDW755 *gmk*<sup>Y163D</sup> | This work |
| JDW1847 | JDW755 *pyk*<sup>H37R</sup> | This work |
| JDW1843 | JDW755 *pyk*<sup>D66N</sup> | This work |
| JDW1838 | JDW755 *pyk*<sup>D125A</sup> | This work |
| JDW1840 | JDW755 *pyk*<sup>A190V</sup> | This work |
| JDW1857 | JDW755 *pyk*<sup>G318E</sup> | This work |
| JDW1832 | JDW755 *pyk* +423 Nonsense | This work |
| JDW1846 | JDW755 *pyk* +468 A insertion | This work |
| JDW1830 | JDW755 *pyk* +783 A insertion | This work |
| DK847 | NCIB3610 pBS32<sup>-</sup> | Yang et al. (49) |
| JDW2231 | DK847 $\Delta yjbM \Delta ywaC relA::mls$ | This work |
| JDW2234 | DK847 $\Delta pyk$ | This work |
| JDW2235 | JDW2231 $\Delta pyk$ | This work |
| DK3287 | NCIB3610 $\Delta zpdN \Delta SP\beta \Delta PBSX \Delta comI$ | Myagmarjav et al. (50) |
| JDW3420 | DK3287 $\Delta pyk$ | She et al. (35) |
| JDW3462 | DK3287 *ndk::Kan* | This work |
| JDW4971 | DK3287 $\Delta pyk, ndk::Kan$ | This work |

Efficient GTP production by pyruvate kinase is not unique to *B. subtilis*. We found that *in vitro*, pyruvate kinases from the facultative anaerobic pathogens *Enterococcus faecalis* and *Listeria monocytogenes*, also showed highest catalytic efficiencies on GDP (Fig. 6F and G). *E. faecalis* PK is also able to use ADP with high efficiency and use UDP efficiently. *L. monocytogenes* PK can use ADP and UDP but less efficiently than GDP. In all three species, CDP is the least efficiently used substrate of pyruvate kinase. These results suggest that in *B. subtilis* and other Firmicutes, GTP can be efficiently produced when a mixture of NDPs is available for substrate-level phosphorylation by pyruvate kinase during glycolysis.

### Pyruvate kinase predominantly generates GTP and UTP as primary nucleotide products during glycolytic growth

Collectively, our findings indicate that pyruvate kinase efficiently produces GTP in (p)ppGpp^0^ cells during guanosine treatment, contributing to guanosine toxicity. However, it remains unclear if pyruvate kinase primarily produces GTP in wild-type cells. Because wild-type cells produce (p)ppGpp, we initially investigated whether (p)ppGpp regulates pyruvate kinase (PK) activity in GTP synthesis, similarly to its inhibitory effects on Gmk and HprT. However, our results indicated little to no effect of (p)ppGpp on PK activity, regardless of whether GDP or ADP was used as the substrate (Fig. 6H).

Next, we performed *in vitro* enzymatic assays with a mixture of ADP and GDP at their physiological concentrations during homeostatic growth (estimated from (33)) of wild-type *B. subtilis*. Under these conditions, ADP concentrations were higher than GDP (0.5 mM ADP vs. 0.2 mM GDP). We monitored ATP and GTP production over time, using anion exchange chromatography that separated these nucleotides to distinct fractions (Fig. S5A). Despite the lower GDP concentration, pyruvate kinase produced both ATP and GTP simultaneously, with GTP being the predominant product (Fig 6I and J). This suggests that GTP, rather than ATP, is the main nucleotide produced by pyruvate kinase.

### Pyruvate kinase and Ndk are the major contributors to NTP synthesis and complement each other during glycolytic growth

Finally, we quantified the levels of NTPs and NDPs, as well as NTP/NDP ratios, in wild-type (WT) and a pyruvate kinase deletion strain (Δ*pyk*) during steady-state glycolytic growth in the absence of exogenous guanosine (Fig. 7A, and Fig. S6). We observed a much higher NTP/NDP ratios for all four nucleotides in wild-type compared to Δ*pyk* mutants, highlighting the strong contribution of PK to NTP synthesis. Among which, GTP/GDP stands as the highest ratio, supporting the hypothesis that pyruvate kinase is a major producer of GTP during glycolytic growth.

**FIG 7.**
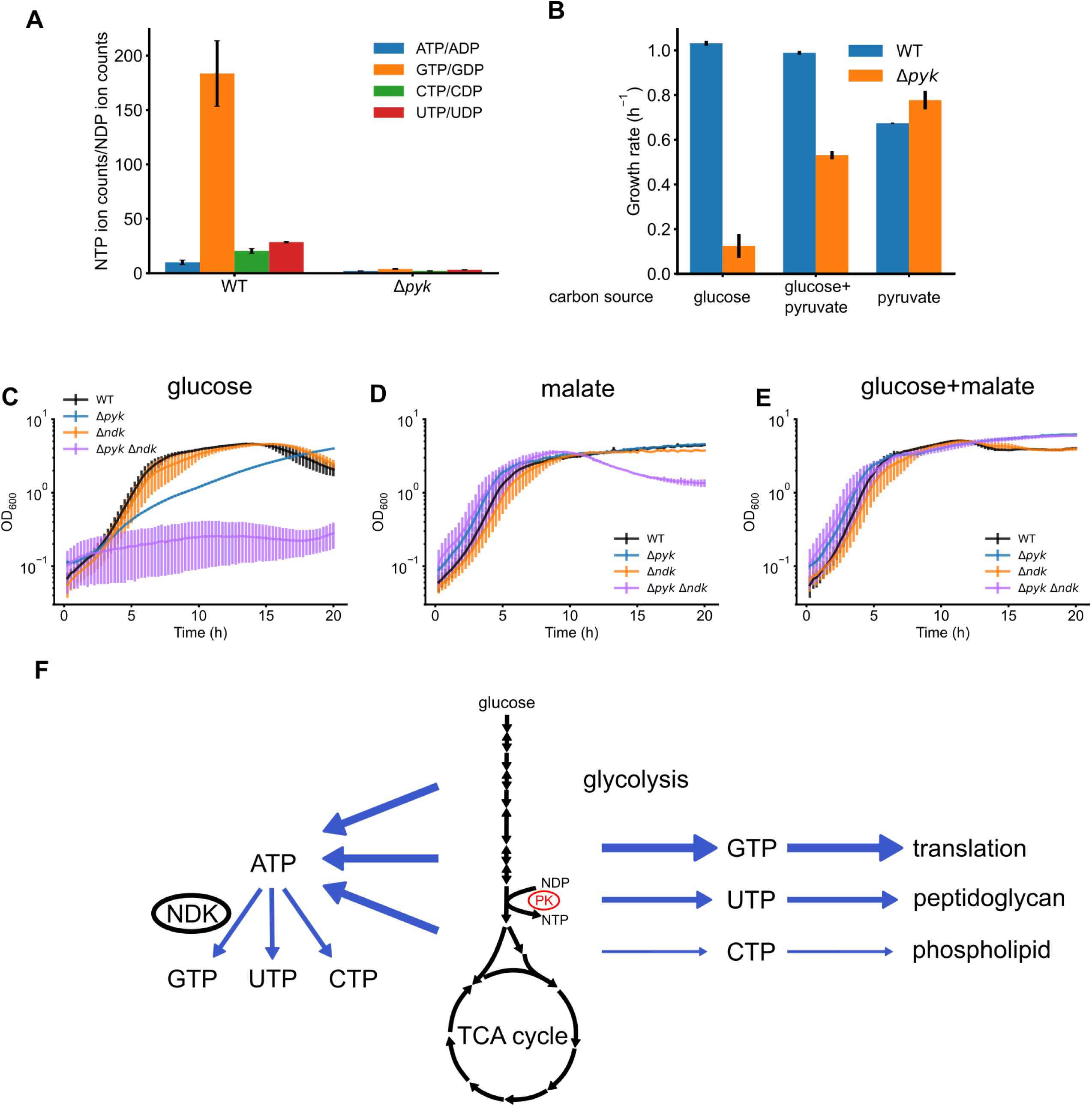
Physiological roles of pyruvate kinase in NTP production and glycolytic growth of *B. subtilis* cells. (A) NTP/NDP ratio in wild-type and Δ*pyk* cells during steady-state growth, metabolites were measured by LC-MS and normalized over OD_600_. (B) Growth rates of WT and Δ*pyk* cells in defined minimal media supplemented with glucose, glucose+pyruvate, or pyruvate, respectively. (C-E) Growth curves of WT, Δ*pyk*, Δ*ndk* and Δ*pyk* Δ*ndk* cells in media with glucose (C), malate (D) or glucose+malate (E). (F) Schematics of NTP synthesis from central carbon metabolism. Pyruvate kinase produces NTPs that directly fuel macromolecular biosynthesis, instead of NTPs produced by Ndk from ATP.

We found that the Δ*pyk* mutant displayed severe growth defects when glucose was used as the sole carbon source. Supplementing the glucose medium with pyruvate, the carbon product of pyruvate kinase (PK), partially alleviated the growth defects (Fig. 7B). The incomplete rescue may be attributed to reduced nucleotide synthesis in the absence of PK activity, or to catabolic repression of the pyruvate transporter (34). Notably, the Δ*pyk* mutant grows well on pyruvate alone (Fig. 7B), consistent with previous findings that *B. subtilis* efficiently imports and utilizes pyruvate in the absence of glucose (35).

To precisely assess the role of pyruvate kinase in nucleotide production, we created a double knockout mutant of *pyk* and *ndk*, a non-essential gene (36, 37) which encodes nucleoside diphosphate kinase that catalyzes the conversion of nucleoside diphosphates (NDPs) to nucleoside triphosphates (NTPs), and assessed its growth. Triple mutant of *ndk* and *pykA pykF* in *E. coli* is viable (19). Strikingly, the *B. subtilis* Δ*ndk*Δ*pyk* mutant failed to form colonies on solid media with glucose as the sole carbon source and showed little growth in liquid glucose media (Fig 7C). By contrasts, the double mutant grew exceptionally well with the gluconeogenic carbon source malate, or both glucose and malate (Fig 7D and E). Since Ndk’s only known function is to synthesize NTPs, the synthetic lethality observed in the Δ*ndk* Δ*pyk* mutant indicates that pyruvate kinase is the major source of NTPs during glycolytic growth in addition to Ndk. In the absence of both enzymes, *B. subtilis* cannot produce sufficient NTPs from glycolysis to sustain growth.

## Discussion

GTP is an essential and abundant nucleotide, serving both as a substrate for transcription and as the energy source for translation. In Gram-positive bacteria, GTP levels are tightly regulated by the alarmone (p)ppGpp, and dysregulation can result in GTP toxicity with exogenous guanosine (13, 17, 18). In this study, we conducted a suppressor screen for guanosine toxicity in a (p)ppGpp^0^ background and identified loss-of-function mutations that led to reduced toxicity and restored growth. The suppressor mutations were mapped to three genes*: hprT*, *gmk*, and *pyk*. The first two genes, *hprT* and *gmk*, are involved in purine salvage and GDP synthesis, and their inactivation reduced GMP and GDP production, respectively (22, 38). The third gene, *pyk*, encoding pyruvate kinase, is a glycolytic enzyme that we surprisingly found to predominantly synthesize GTP instead of its established product ATP. Inactivation of pyruvate kinase lowered GTP production, leading to growth rescue in the presence of guanosine. Our findings also suggest that the unexpected GTP synthesis activity of pyruvate kinase may be necessary to fulfill the high cellular GTP demand for active transcription and translation in wild type cells.

Our suppressor screen identified loss-of-function mutations in *pyk* (encoding pyruvate kinase) that mitigate guanosine toxicity by reducing GTP production from GDP. Metabolomic analyses revealed that the sharp increase in GTP levels associated with guanosine toxicity is partially alleviated in *pyk* suppressors. Notably, no suppressor mutations were found in *ndk*, a non-essential gene (36, 37), which encodes nucleoside diphosphate kinase that catalyzes the conversion of nucleoside diphosphates (NDPs) to nucleoside triphosphates (NTPs), including GDP to GTP. This leads to a hypothesis that pyruvate kinase not only produces GTP more efficiently than ATP or other NTPs but may serve as the primary source of GTP synthesis during glycolysis in *Bacillus subtilis*.

In addition to nucleotide changes, we observed higher levels of glycolytic intermediates upstream of pyruvate kinase in the (p)ppGpp^0^ *pyk* suppressor compared to (p)ppGpp^0^. This result is expected because pyruvate kinase converts the lower glycolytic intermediate phosphoenolpyruvate (PEP) to pyruvate, depleting PEP and its upstream intermediates. Loss-of-function mutations in pyruvate kinase halt the consumption of PEP, leading to its accumulation. Interestingly, not all glycolytic intermediates upstream of pyruvate kinase respond similarly across the mutant backgrounds (Fig. 4B). Reactions between PEP and 1,3-bisphosphoglycerate (BPG) are physiologically reversible, resulting in a consistent behavior of metabolites in this segment, with 8–10-fold higher levels in (p)ppGpp^0^ *pyk* compared to (p)ppGpp^0^. However, the conversion of glyceraldehyde 3-phosphate (G3P) to BPG, catalyzed by the glycolytic enzyme GapA in *B. subtilis*, is irreversible. The other glyceraldehyde 3-phosphate dehydrogenase (GAPDH), GapB, is not expressed during glycolysis, making GapA the sole enzyme available for this step. Consequently, the accumulation of BPG does not significantly affect levels of glycolytic intermediates upstream of G3P, such as DHAP, G6P and FBP. As a result, glycolytic intermediates upstream of GapA show only modest (∼2-fold) differences.

Our discovery that pyruvate kinase selectively produces GTP and UTP during glycolysis revealed a novel connection between carbohydrate metabolism and nucleotide synthesis (Fig. 7F). These results challenge the conventional view of nucleotide synthesis through glycolysis, which emphasizes that ATP is the main energy nucleotide produced by both phosphoglycerate kinase (PGK) and pyruvate kinase, with other NTPs synthesized from NDPs via ATP donation (4). Our data show that, at least in *B. subtilis* and related Firmicutes, GTP is directly synthesized from GDP during glycolysis, bypassing the need of Ndk using ATP as an intermediate. Additionally, UTP is also a significant product of pyruvate kinase, indicating that it can be directly synthesized from glycolysis. The direct synthesis of GTP and UTP likely provides a significant advantage, as the efficient production of all four ribonucleotides (ATP, GTP, UTP, and CTP) is essential for bacterial growth and macromolecular synthesis. GTP powers protein synthesis directly, while UTP initiates peptidoglycan synthesis through UDP-GlcNAc production (39). The nucleotide selectivity of pyruvate kinase may be specifically tailored to meet the cellular demand for these different macromolecules, rather than relying on indirect production via Ndk. This would reduce the cellular burden on Ndk and metabolic flux, ensuring optimal resource allocation and supporting efficient growth. The synthetic lethality of the Δ*ndk* Δ*pyk* double mutant specifically on glucose suggests that these two enzymes complement each other in producing the complete set of (d)NTPs required for optimal growth under glycolytic conditions.

The selectivity of PK to produce various NTPs varies for different species. For example, pyruvate kinases from Firmicutes, including *B. subtilis* (Fig. 6E), *Listeria* (Fig. 6G) and *S. pneumoniae* (40), favor GTP production, *E. faecalis* PK produces both ATP and GTP efficiently (Fig. 6F). In *E. coli* there are two pyruvate kinase isoforms, PK I (encoded by *pykF*) and PK II (encoded by *pykA*). *In vitro*, PK I appears to favor GTP and PK II favors ATP (30, 31). While both genes are required for optimal growth, explanation was only provided in terms of their roles in pyruvate synthesis (41). Little is known about the relative contributions of the two isoforms during glycolytic growth to GTP synthesis.

It is noteworthy that the Δ*ndk*Δ*pyk* double mutant remains viable on gluconeogenic carbon sources, suggesting another enzyme(s) is producing NTPs, likely through the TCA cycle. Substrate level phosphorylation in the TCA cycle, through succinyl-CoA synthetase, has been known to produce GTP in certain species. In *E. coli*, succinyl-CoA synthetase preferentially makes ATP; in *Thermus aquaticus,* succinyl-CoA synthetase favors GTP synthesis (42). In mitochondria, succinyl-CoA synthetase has evolved two homologs—one that preferably produces GTP and the other that synthesizes ATP (43, 44). This specificity may reflect an evolutionary adaptation to optimize nucleotide production in different bacterial species and mitochondria.

Despite our unexpected findings on *B. subtilis* pyruvate kinase’s preference for GTP over ATP, the exact cause of “death-by-GTP” remains unresolved. Initially, we hypothesized that guanosine toxicity in (p)ppGpp^0^ cells was due to the depletion of key metabolites like G6P, leading to misallocation of resources and deficiencies in cell envelope or pentose phosphate pathway precursors, which was alleviated in *pyk* suppressors. However, our current data do not support this hypothesis. While we recently showed that glycolytic intermediate expansion due to inactivation of pyruvate kinase is important for efficient and robust gluconeogenesis, it does not appear to benefit steady state growth during glycolysis (35). Similarly, in this work, none of the observed changes in glycolytic intermediates explain guanosine toxicity. The source of GTP toxicity thus remains an open question.

Although it remains mechanistically enigmatic, GTP toxicity may not be confined to bacteria, as elevated GTP levels generated through substrate-level phosphorylation in the TCA cycle within mitochondria have been linked to reproductive aging. For instance, in *C. elegans* mitochondria, the GTP-producing succinyl-CoA synthetase has been associated with reproductive aging, whereas its ATP-producing counterpart does not show this effect (44). While direct GTP synthesis by carbon metabolism enzymes can enhance metabolic efficiency, it also introduces vulnerabilities, such as an increased risk of DNA damage. Further research will be critical to addressing these questions and advancing our understanding of the role and consequences of direct GTP synthesis by substrate-level phosphorylation in both bacterial and mitochondrial metabolism.

## Materials and Methods

### Strains and growth conditions

All *Bacillus subtilis* strains were derived from YB866 or NCIB3610 (Table 2). *B. subtilis* was grown in S7 defined liquid medium (45) with MOPS used at 50 mM instead of 100 mM, or solid medium with Spizizen salts (Spiz), supplemented with 1% (w/v) glucose, 0.5% (w/v) casamino acids (CAS) and 40 µg ml^−1^ tryptophan and methionine (13) when necessary. Guanosine was suspended in sterile Milli-Q water and used at a final concentration of 0.1 mM in agar plates and 1 mM in liquid broth.

### Suppressor selection and verification

Single colonies of (p)ppGpp^0^ were grown independently in S7 + CAS medium to saturation. An aliquot of 100 µl culture was plated directly onto Spiz + CAS with 0.1 mM guanosine. Only one suppressor was selected from each plate and the suppressor was verified by its ability to grow on Spiz + CAS agar plates supplemented with 0.1 mM guanosine.

### Suppressor mutation identification

Sanger sequencing of target genes in the GTP biosynthesis pathway was performed for the low GTP group (≤ 2 fold change of GTP levels relative to untreated). Mutations in the intermediate GTP group (>2 fold change of GTP levels relative to untreated) were identified by whole genome sequencing and Sanger sequencing of target genes as described previously (13, 46).

### Thin layer chromatography (TLC)

TLC was performed exactly as described(15). Briefly, *B. subtilis* cells were grown in low phosphate (0.5 mM) S7 media with 0.5% (w/v) casamino acids and labeled with 50 µCi ml^−1^ ^32^P orthophosphate (900 mCi mmol^−1^; Perkin-Elmer) for 2 – 3 generations before sampling. Nucleotides were extracted by incubating 100 µl culture with 20 µl of 2 M formic acid on ice for at least 20 minutes. Samples were spotted onto a cellulose PEI plate and separated in 1.5 M KH_2_PO_4_ (pH 3.4). The TLC plates were exposed to a storage phosphor screen. Nucleotides were detected by scanning with a Typhoon scanner (GE) and quantified using the ImageQuant software (Molecular Dynamics).

### LC-MS metabolomics

Cells were grown in modified S7 media with 5 mM MOPS until OD_600_ reached ∼0.3. When required, guanosine was added to a final concentration of 1 mM. Before and after guanosine treatment, 5 ml samples were taken for metabolite extraction and LC-MS analysis as described previously (21).

### Expression, purification, and enzymatic assay of pyruvate kinase

The coding sequence of *B. subtilis pyk* was PCR amplified and cloned into the pLICtrPC-HA vector by ligation independent cloning (47). The recombinant plasmid was verified by Sanger sequencing and transformed into *Escherichia coli* BL21 (DE3) cells (Agilent Technologies). Transformants were grown at 37 °C in Lysogeny Broth (LB) with 100 μg ml^−1^ carbenicillin to an OD_600_ ∼ 0.5 and induced with 1 mM IPTG for 3 hours before harvest. Cell pellets were disrupted in 300 mM NaCl, 25 mM Tris-HCl (pH 7.5), and 10 mM imidazole by passing through a French press twice. Cell debris was cleared by centrifugation and the supernatant was loaded onto a HisTrap FF column (GE Healthcare) using an ÄKTA FPLC (GE Healthcare). The N-terminally His-tagged pyruvate kinase was eluted using a linear gradient of imidazole in 300 mM NaCl and 25 mM Tris-HCl (pH 7.5) and dialyzed in 40 mM Tris-HCl (pH 7.5), 100 mM NaCl, 0.5 mM EDTA, and 1 mM DTT. The His-tag was cleaved by a His-tagged TEV protease (1 mg TEV protease for 1L culture), and untagged pyruvate kinase was collected by flowing through HisTrap FF column. Purified pyruvate kinase was analyzed by SDS-PAGE and quantified by Bradford assay (Bio-Rad).

Activity of pyruvate kinase was assayed at 25°C in a 100 μL mix containing 100 mM Tris-HCl (pH 7.5), 100 mM KCl, 10 mM MgCl_2_, 1.5 mM phospho(enol)pyruvate (PEP), 150 μM NADH, and 2U L-lactic dehydrogenase (from bovine muscle, Sigma-Aldrich), various concentration of ADP or GDP, and 20 nM pyruvate kinase. Reactions were monitored by measuring decrease of A_340_ in a Shimadzu UV-2401PC spectrophotometer for up to 5 minutes.

### ADP/GDP competition assay with pyruvate kinase

Reaction was prepared in a 400 μL mixture containing 100 mM Tris-HCl (pH 7.5), 100 mM KCl, 10 mM MgCl_2_, 200 μM PEP, 500 μM ADP, 200 μM of GDP, 100 μM AMP, and 20 nM pyruvate kinase. The reaction was periodically examined by mixing 350 μL of sample with 14 μL of 0.5 M EDTA at 1 min, 2 min, 3 min, 5 min and 10 min to stop the reaction. 364 μL mix was diluted to 5 ml with ddH_2_O, and then loaded onto a HiTrap QFF 1 ml column (GE Healthcare) using an ÄKTA FPLC (GE Healthcare). GTP and ATP were eluted using a NaCl gradient in 25 mM citric acid, pH 3.5, as described below. Samples were loaded and thoroughly washed in 25 mM citric acid (pH 3.5), followed by elution by a linear gradient of 0 mM – 60 mM NaCl (5 column volumes (CVs)), a constant gradient of 60 mM NaCl (10 CVs), and another linear gradient of 60 mM – 300 mM NaCl (15 CVs). Absorbance at 254 nm was recorded to detect nucleotides. AMP and ADP were eluted before 60 mM NaCl, GDP was eluted at ∼60 mM NaCl, ATP and GTP were eluted between 60 mM – 300 mM NaCl. The amounts of ATP and GTP were quantified using pure standards of known quantities.

## Acknowledgements

We thank Petra Levin for helpful discussions. This work is supported by the USA National Institute of Health R35 GM127088 (to J.D.W.), the National Science Foundation (NSF) grant award no. 1715710 (to D.A.-N.). B.W.A. was supported by NSF GRFP DGE-1256259.

